# Theoretical design of a space bioprocessing system to produce recombinant proteins

**DOI:** 10.1101/2022.09.20.508767

**Authors:** Mathangi Soundararajan, Matthew B. Paddock, Michael Dougherty, Harry W. Jones, John A. Hogan, Frances M. Donovan, Jonathan M. Galazka, A. Mark Settles

**Affiliations:** KBR, NASA Ames Research Center, Moffett Field, CA 94035; Bioengineering Branch, NASA Ames Research Center, Moffett Field, CA 94035; Space Biosciences Research Branch, NASA Ames Research Center, Moffett Field, CA 94035

## Abstract

Space-based biomanufacturing has the potential to improve the sustainability of deep space exploration. To advance biomanufacturing, bioprocessing systems need to be developed for space applications. Here, commercial technologies were assessed to design space bioprocessing systems to supply a liquid amine carbon dioxide scrubber with active carbonic anhydrase produced recombinantly. Design workflows encompassed biomass dewatering of 1 L *Escherichia coli* cultures through to recombinant protein purification. Equivalent system mass (ESM) analyses had limited utility for selecting specific technologies. Instead, bioprocessing system designs focused on minimizing complexity and enabling system versatility. Three designs that differed in biomass dewatering and protein purification approaches had nearly equivalent ESM of 357-522 kg eq. Values from the system complexity metric (SCM), technology readiness level (TRL), and degree of crew assistance metric identified a simpler, less costly, and easier to operate design for automated biomass dewatering, cell lysis, and protein affinity purification.

## Introduction

Liquid amine scrubbing is a promising technology to capture CO_2_ produced during crewed extraterrestrial missions^1,2^. In this system, a CO_2_ -rich gas stream is passed over an organic liquid amine which absorbs the CO_2_. Pure CO_2_ is recovered by raising the solution temperature, which regenerates the amine for subsequent capture at low temperature^3^. An efficient liquid amine system should have fast absorption kinetics to reduce system size as well as a low desorption temperature to minimize energy inputs. Different liquid amines either have fast absorption kinetics or low desorption temperature but not both simultaneously^4^. Carbonic anhydrase is an enzyme that catalyzes the interconversion of CO_2_ and HCO_3_ ^-^ to increase the CO_2_ sequestration in some liquid amine systems. Addition of carbonic anhydrase enhances the absorption kinetics of liquid amines with a low desorption temperature enabling both reduced size and improved energy efficiency of CO_2_ scrubbing^5–7^.

Implementing an enzyme-assisted liquid amine CO_2_ scrubber requires a time-course supply of carbonic anhydrase. On long-duration space missions close to Earth, this requirement could be met by resupply or long-term storage. However, resupply is not an economical option for many deep space missions like those planned for Mars. Purified enzymes are typically sensitive to room temperature conditions and require ultra low storage temperatures to retain activity long-term. Not all enzymes retain activity in low temperature storage and in situ production of enzymes would mitigate risks of relying solely on low temperature storage of proteins with limited stability. Space biomanufacturing systems have the potential to produce enzymes and other biological materials using in situ resources during a Mars mission^8–13^. Space systems must minimize cost and crew time, while assuring astronaut safety and addressing effects of increased radiation and reduced gravity^14–17^.

Previous space biomanufacturing studies and reviews evaluated large-scale mission design^8,18,19^, microbial growth kinetics^20^, and bioreactor design^20–22^. Extracting products of interest at sufficient quality is equally essential to develop biomanufacturing. Systems such as Wetlab-2 or the Gene Expression Measurement Module (GEMM) illustrate the challenge of adapting biological sample processing for RNA extraction and molecular analysis in the microgravity environment^23,24^. In these systems, the sample mass and volume processed was small with the goal of providing biological inputs for analytical experiments^23,24^. Future space biomanufacturing systems need to address post-growth processing at larger scales to extract products for in situ use.

In this study, we compared commercial technologies and potential designs for in space biomanufacturing systems. Our operational scenario was post-growth bioprocessing to produce recombinant carbonic anhydrase from *Escherichia coli* during a Mars mission. Since carbonic anhydrase is unlikely to be the only useful product in deep space missions, the ability to produce a variety of recombinant proteins from multiple chassis organisms was a key consideration for the designs. The designs were compared using equivalent system mass (ESM) analysis^14,25^, a system complexity metric (SCM)^16^, technology readiness level (TRL)^26^, and crew-mediated steps to guide future prototype development efforts.

## Results

Bioprocessing technology comparisons were based on a production scenario that required thermostable and high-pH tolerant carbonic anhydrase for a liquid amine CO_2_ capture system during 600 days of surface operations on Mars^27,28^. In this system, enzyme activity will decay with heating-cooling cycles, and will require intermittent addition of recombinant protein to supply sufficient enzyme activity throughout the surface operations. Protein purification was assumed to be required to reduce side reactions with the liquid amine; however, multiple chromatography steps to produce highly purified protein are not required for this application^5^. Active enzyme can be produced in *E. coli* as an intracellular recombinant protein with a His-tag for affinity purification. A prior study reported 180 mg/L of recombinant carbonic anhydrase yield^29^. A total of 14.4 g of active enzyme was estimated to be required to supply a crew of six for 600 days, and 100 cultures at 1 L volume is estimated to be sufficient.

### Potential workflows for the operational scenario

Figure 1 shows five potential bioprocessing workflows (a, b, c, d, and e) starting from a common cell growth and production step. Each workflow considers sub-processes of biomass processing, protein extraction, and storage. These sub-processes were split into steps that could have multiple alternate methods. For the sub-process of biomass processing, dewatering and drying steps were the primary options considered. The stated use case will produce recombinant protein intracellularly and will require protein extraction sub-process including cell lysis, protein purification, and buffer exchange/desalting steps. Finally, protein product could have a storage sub-process, either as biomass or as a purified product.

**Fig. 1.**
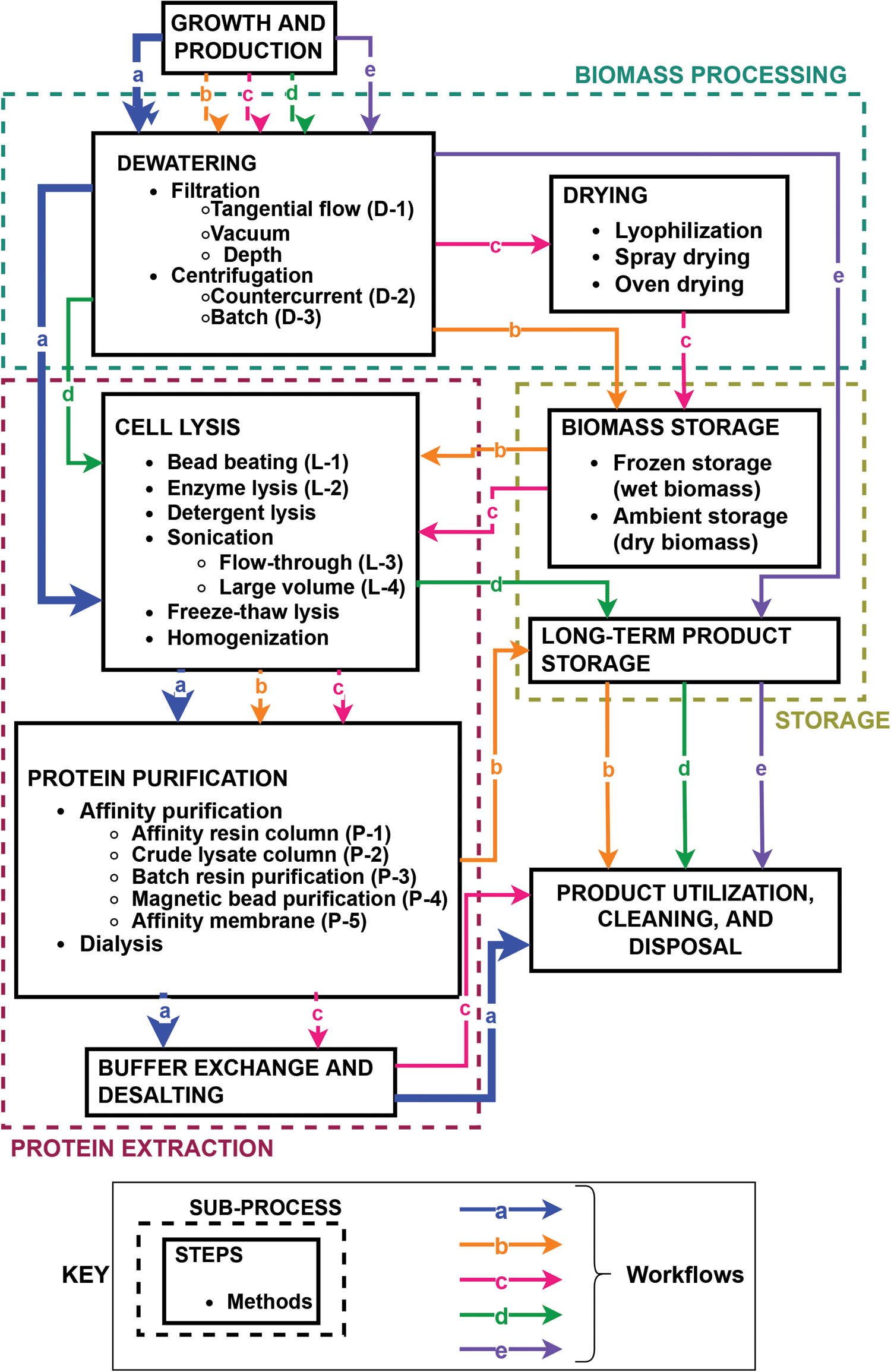
Flow diagram of potential bioprocessing strategies. Dashed boxes group the primary sub-processes of biomass processing, protein extraction, and product storage. Solid boxes give individual steps with bulleted lists of common methods to complete the step. Methods compared in this analysis have parenthetical designations that are used in Fig. 2 and Fig. 3. Lettered arrows (a, b, c, d, e) give examples of possible workflows for processing E. coli cells expressing recombinant carbonic anhydrase.

Workflow-a was selected as best aligned with the operational scenario of supplying purified carbonic anhydrase on a 6 to 8-day cycle. This workflow moves from growth and production steps to dewatering, cell lysis, protein purification, and buffer exchange to end with product utilization, cleaning, and disposal. Although storage and drying steps were considered, they were not included and deemed unnecessary based on the selected scenario.

### Biomass processing

Dewatering cultures greatly reduces processing volumes for protein extraction or biomass storage sub-processes. However, it may be feasible to complete protein extraction without a dewatering step. We modeled the impact of dewatering on cell lysis and protein purification steps by scaling processing time for flow-through methods including bead beater lysis (L-1), flow cell sonication (L-3), affinity resin column purification (P-1), and crude lysate column purification (P-2) methods (Fig. 2, Supplementary data 1). This analysis showed that processing cultures without dewatering would require multiple days using flow-through methods.

**Fig. 2.**
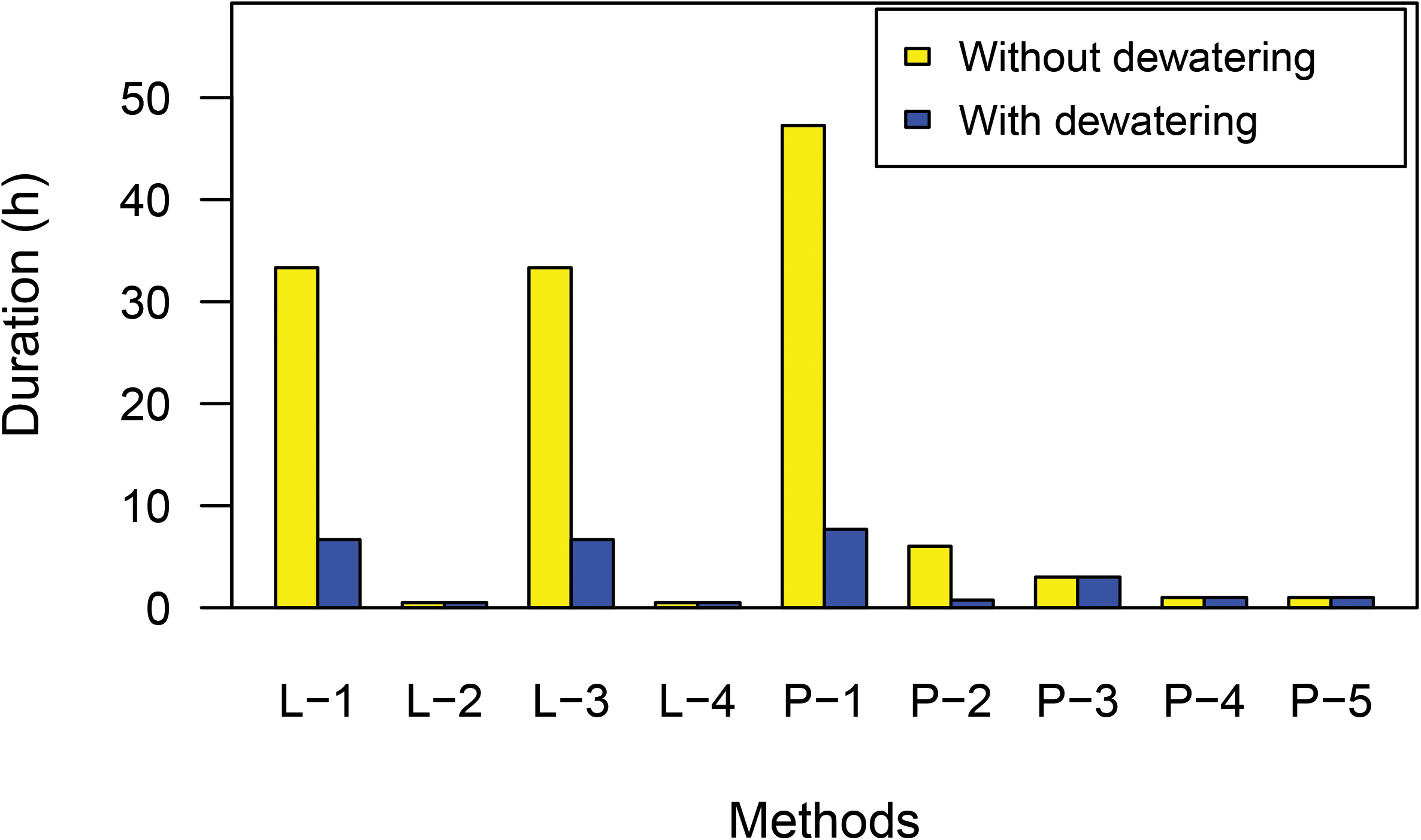
Estimated process duration for each run of the lysis (L) and purification (P) based on recommended manufacturer protocols or published literature. Methods included: flow through bead beater (L-1), enzyme lysis (L-2), flow cell sonicator (L-3), large volume probe sonicator (L-4), affinity resin column (P-1), crude lysate column (P-2), batch affinity resin (P-3), batch affinity magnetic beads (P-4), and affinity membrane (P-5). Black bars are process times of 1 L culture and gray bars are process times of 200 mL concentrated biomass. See supplementary data 1 for detailed calculations.

Other methods could be completed in batch including enzyme lysis (L-2), large-volume probe sonication (L-4), batch affinity resin purification (P-3), magnetic bead affinity purification (P-4), and affinity membrane purification (P-5). These were scaled by increasing materials to limit impacts on processing duration (Fig. 2). However, most protein purification methods were developed for cell lysates from concentrated biomass suspensions. It is unknown whether cultures could be lysed directly, or if large-scale batch purification would yield sufficiently concentrated purified protein. We concluded that dewatering would help ensure feasibility for batch methods and allow flow methods to be considered as options in the bioprocessing designs.

ESM is a metric to estimate the flown mass required to implement a space technology and can help guide selection of alternative technologies^14,25,30^. The metric is a linear model to estimate mass of the system along with mass equivalences for providing volume and other infrastructure within a spacecraft (Supplementary data 1). Table 1 gives the 2015 and 2022 equivalency factors used to calculate ESM ^31,32^, and Table 2 reports total ESM for the methods and bioprocessing designs. These calculations were highly correlated (r = 0.992), and the 2015 equivalency factors are reported in the figures to enable comparisons to prior technology proposals. Due to the variability in calculating ESM for preliminary designs, the metric is only used for selection of competing technologies when ESM estimates approach a 5-to 10-fold difference^25,33^. Figure 3a shows ESM calculations for dewatering methods. Batch and countercurrent centrifugation have ∼5-fold higher ESM than a tangential flow filter. Despite the similar ESM, a countercurrent centrifuge is easier to automate and can potentially reduce crew time requirements. Based on these considerations, we removed batch centrifugation from further consideration.

**Table 1.** Mars surface equivalency factors for this study

| Parameter | 2015 Equivalency factor | 2022 Equivalency factor |
| --- | --- | --- |
| Shielded Volume (V) | 215.5 kg<sub>eq</sub>/m<sup>3</sup> | 79.3 kg<sub>eq</sub>/m<sup>3</sup> |
| Power (P) | 87 kg<sub>eq</sub>/kW | 162 kg<sub>eq</sub>/kW |
| Thermal Control (C) | 146 kg<sub>eq</sub>/kW | 96 kg<sub>eq</sub>/kW |
| Cold Storage (CS) | 0.79 kg<sub>eq</sub>/kg | 0.21 kg<sub>eq</sub>/kg |
| Water Treatment (W) | 0.12 kg<sub>eq</sub>/kg | 0.3 kg<sub>eq</sub>/kg |
| Waste Storage (WS) | 0.83 kg<sub>eq</sub>/kg | 0.93 kg<sub>eq</sub>/kg |

**Table 2.** Total ESM for the methods and designs reported in this study.

| Method/design | 2015 factors | 2022 factors | Designation |
| --- | --- | --- | --- |
| Tangential flow filter | 50 | 40 | D-1 |
| Countercurrent centrifuge | 290 | 220 | D-2 |
| Batch centrifuge | 120 | 90 | D-3 |
| Bead beating | 14.2 | 14.4 | L-1 |
| Enzyme lysis | 4.1 | 2.6 | L-2 |
| Sonicator (flow cell) | 189 | 201 | L-3 |
| Sonicator (large volume probe) | 182 | 197 | L-4 |
| Affinity resin column | 43 | 32 | P-1 |
| Crude lysate column | 16 | 18 | P-2 |
| Batch resin purification | 46 | 66 | P-3 |
| Magnetic beads | 41 | 46 | P-4 |
| Affinity membrane | 44 | 32 | P-5 |
| Design 1 | 460 | 400 |  |
| Design 2 | 360 | 340 |  |
| Design 3 | 520 | 450 |  |

**Fig. 3.**
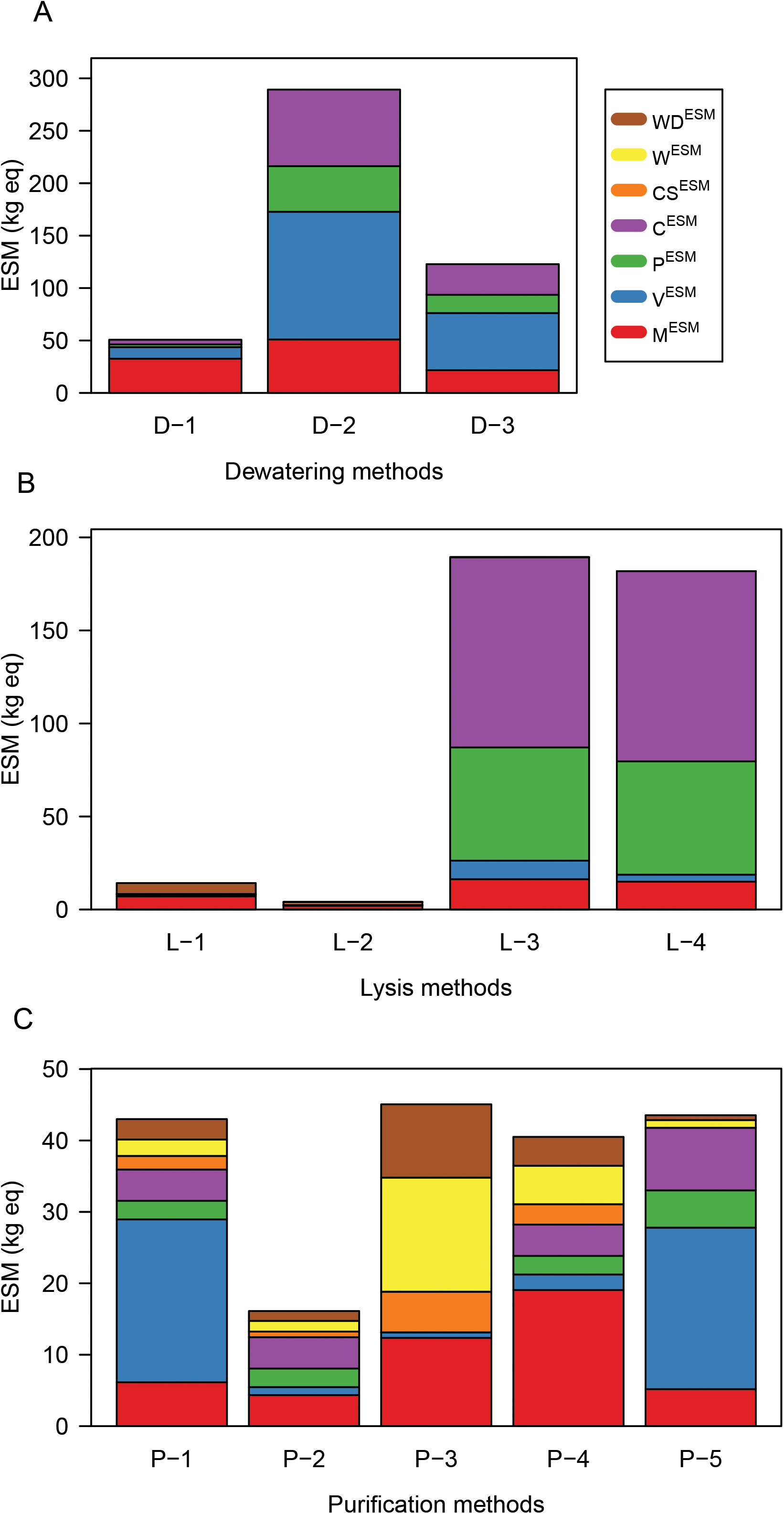
ESM models assuming 100 bioprocessing runs. (a) Dewatering methods included tangential flow filtration (D-1), countercurrent centrifugation (D-2), and batch centrifugation (D-3). (b) Cell lysis methods included bead beating (L-1), enzymatic lysis (L-2), flow through sonication (L-3), and large volume probe sonication (L-4). (c) Protein purification methods included clarified lysate column (P-1), crude lysate column (P-2), batch resin purification (P-3), magnetic bead purification (P-4), and affinity membrane purification (P-5). Each bar graph shows mass equivalencies for: total mass (M^ESM^), volume (V^ESM^), power (P^ESM^), cooling (C^ESM^), cold storage (CS^ESM^), water treatment (W^ESM^), and solid waste disposal (WD^ESM^). See Table 2 for total ESM values and Supplementary data 1 for detailed calculations.

### Protein extraction

ESM models for lysis eliminated sonication due to high power and cooling requirements (Fig. 3b). Although enzyme lysis has a low ESM in this analysis, there are several concerns for implementation of an enzyme-only method that could not be easily modeled via ESM. Lysozyme is only effective as a lysis method for a limited number of microbial host species. The enzyme is most effective when combined with chelators, detergent, or sonication to disrupt the outer membrane of gram-negative bacteria like *E. coli*^34^. The chelators and detergents required for more effective enzyme lysis also create wastewater treatment challenges. Finally, enzyme lysis is not as effective as mechanical lysis^35^ and would likely require greater biomass growth to achieve equivalent recombinant protein yield.

Mechanical lysis with bead beating is a well-established cell disruption technique for a large variety of organisms and developmental stages including spores^36^. A small footprint, flow-through bead beater has been used on the ISS for biology research, demonstrating technology feasiblity^23^. Based on these factors, flow-through bead beating was selected as the lysis method for design comparisons.

ESM estimates for the protein purification step compared five commercial affinity purification methods. ESM estimates were within a 3-fold range for all technologies (Fig. 3c). Like the cell lysis step, we considered feasibility of implementation to select three methods for bioprocessing system design comparisons. The affinity resin column and crude lysate column use equivalent flow-through approaches, but the affinity resin column required a clarified lysate with minimal cell debris. The crude lysate column was selected for increased reliability and reduced complexity. Batch affinity purification using magnetic beads requires a crew-assisted step. By contrast, the batch affinity resin “tea bag” method would be simpler to automate and was selected. The affinity membrane was selected to analyze design requirements for an alternate solid matrix.

### Bioprocessing system integrated designs

Figure 4 shows the steps and methods eliminated and retained to design integrated bioprocessing systems. Table 3 summarizes key decisions and the rationale for selection of specific steps and methods. Biomass storage, product storage, and drying steps were eliminated. For dewatering, the counter current centrifuge and filtration methods were retained. For cell lysis, only a flow-through bead beater was retained. For protein purification, crude lysate column, batch resin purification, and affinity membrane purification were retained.

**Fig. 4.**
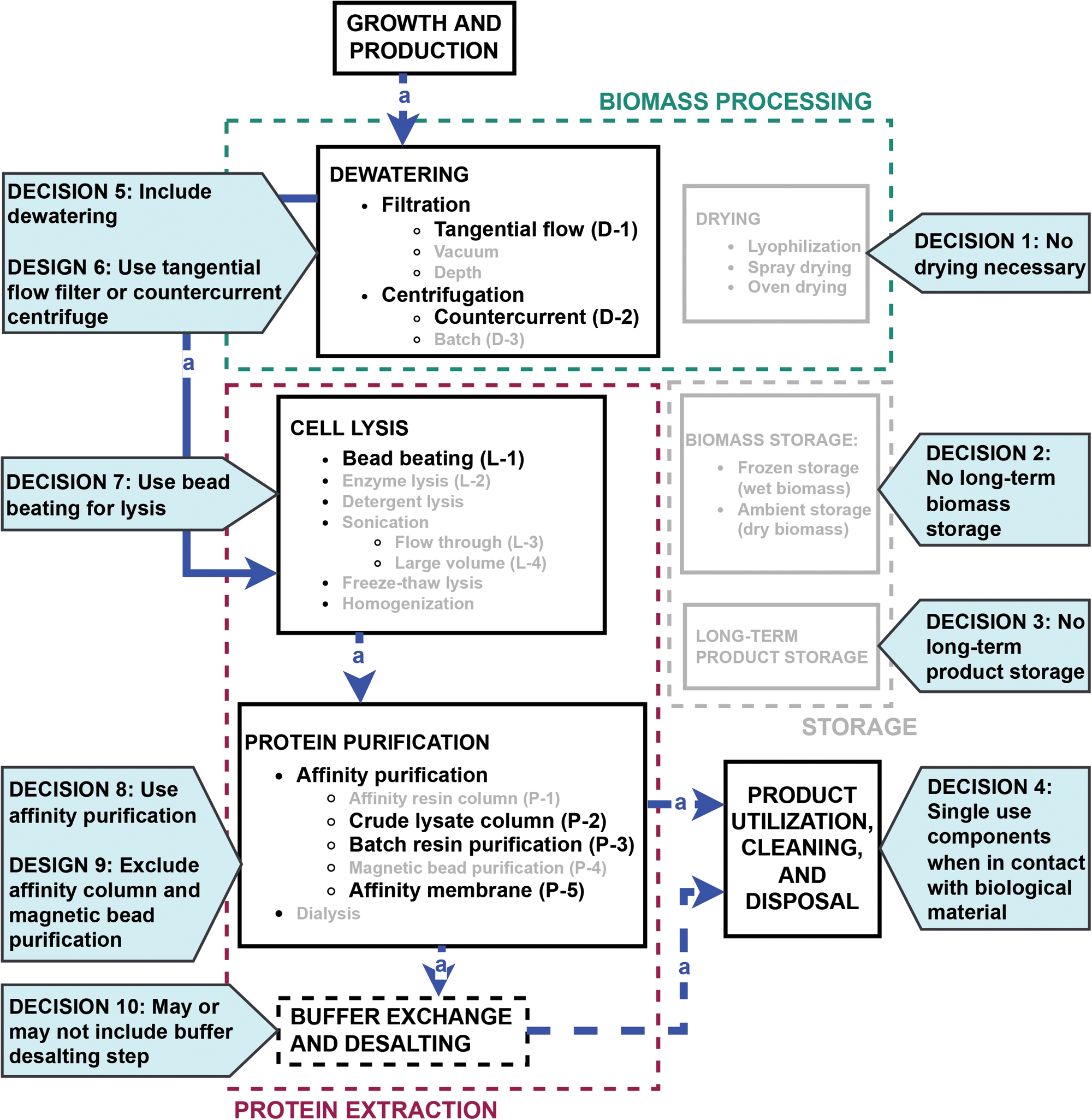
Flow diagram of bioprocessing strategies showing selected methods for integrated, in space designs. Decisions from the trade study are indicated with pink callouts. The rationale for each decision is given in Table 3.

**Table 3.** Summary of workflow method decisions and rationales.

|  | Decision | Rationale | Issues/concerns |
| --- | --- | --- | --- |
| 1 | No long-term biomass storage | • Reduce complexity |  |
| 2 | No long-term product storage | • Reduce complexity | • Increases mission risk in case of anomalies |
| 3 | No drying | • Reduce complexity |  |
| 4 | Any component in contact with biological material will be single use | • Reduce crew time for cleaning<br>• Reduce complexity<br>• Reduce chemical safety concerns |  |
| 5 | Include dewatering | • Shorten processing time | • Adds complexity |
| 6 | Use tangential flow filters or countercurrent centrifuge | • Batch centrifuge requires crew assisted steps | • Countercurrent centrifuge may require modification for microbial applications |
| 7 | Use bead beating for lysis | • Applicable for multiple chassis organisms<br>• Demonstrated in space | • May require modification to process large volumes |
| 8 | Use affinity purification | • Product specific<br>• Produces relatively pure protein after a single purification step | • Must supply affinity binding competitors for elution |
| 9 | Exclude affinity column and magnetic beads | • Crude lysate column may increase reliability<br>• Magnets increase complexity |  |
| 10 | May or may not include a buffer exchange and desalting step | • Insufficient data to either include or eliminate this step |  |

We developed three integrated designs using the methods selected from the trade study (Fig. 5). Design 1 uses a countercurrent centrifugation system to dewater biomass, lyse cells, and purify the protein with a batch resin method. Material flow is mediated by a peristaltic pump (1) and automated pinch valves (2). Cells from the biomass reservoir (3) are fed into the centrifuge (4) and concentrated. Supernatant media is collected in a spent media reservoir (5). Concentrated biomass is pumped through a bead beater (6) for lysis and the lysate is returned to the biomass reservoir. The lysate is then pumped into the affinity resin reservoir (7) for protein binding. Protein-bound resin is separated from the lysate using the centrifuge, while the spent lysate is collected in a waste reservoir (8). The resin is washed with buffer (9). Wash buffer is separated by the centrifuge and collected into the waste reservoir. The bound protein is eluted from the resin using elution buffer (10), separated by the centrifuge, and collected in the product reservoir (11).

**Fig. 5.**
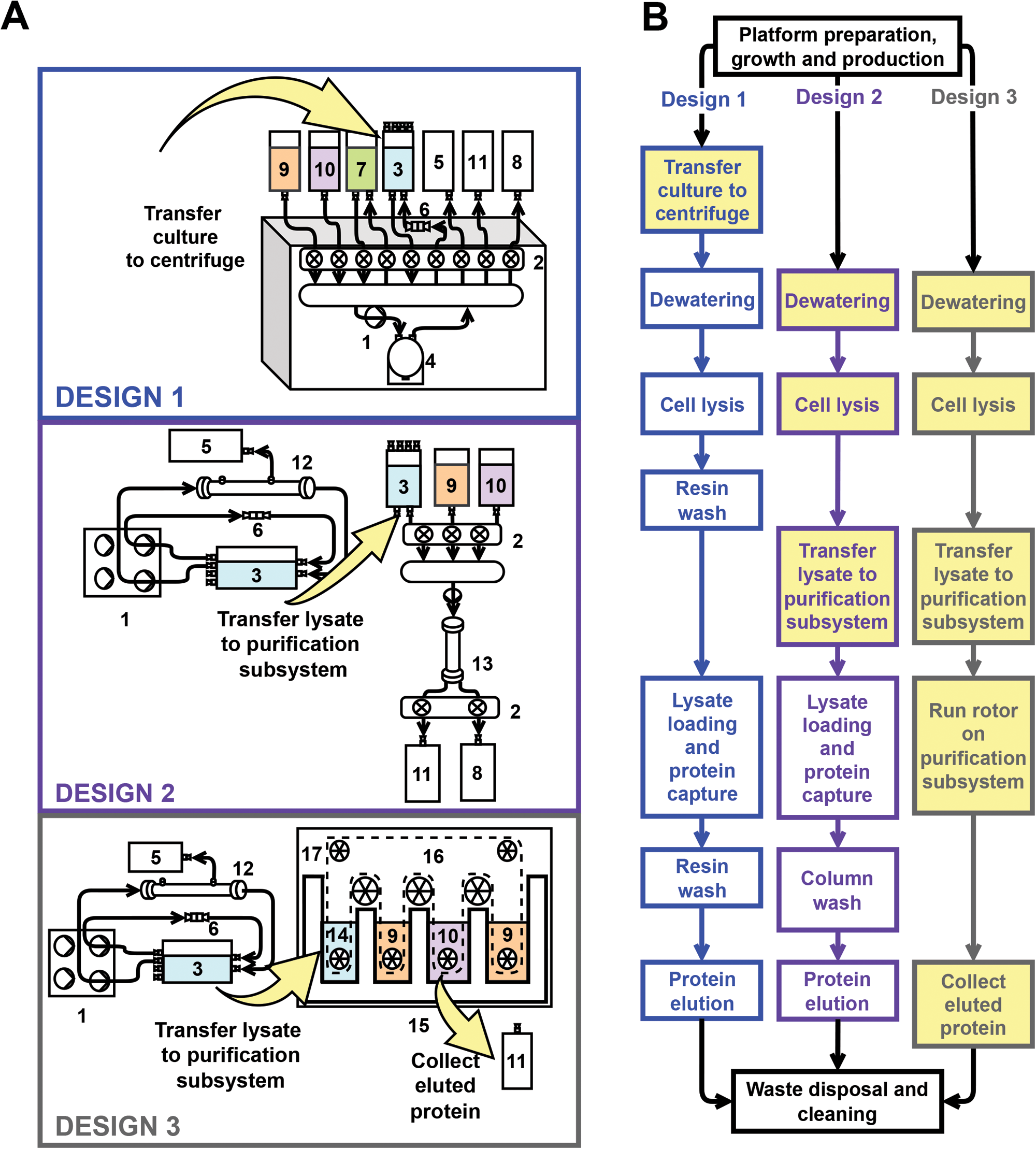
Bioprocessing system designs integrating selected methods for dewatering, lysis, and purification. (a) Design schematics with the following components: peristaltic pump (1), pinch valve (2), biomass reservoir (3), centrifuge cartridge (4), spent media reservoir (5), disposable bead beater (6), affinity resin reservoir (7), waste reservoir (8), wash buffer (9), elution buffer (10), product reservoir (11), tangential flow filter (12), crude lysate column (13), crude lysate chamber (14), affinity purification cartridge (15), affinity membrane (16), rollers (17). Yellow arrows indicate crew-assisted steps. (b) Flow diagram comparing the bioprocessing designs. Yellow boxes indicate methods that require crew-assistance to initiate or complete the method.

Design 2 assumes a peristaltic pump (1) from the growth and production step will pump fluids through a tangential flow filter (12) to dewater and concentrate biomass. Concentrated biomass is returned to the biomass reservoir (3), and clarified media is collected in the spent media reservoir (5). The concentrated biomass is lysed by the bead beater (6), and crude lysate is returned to the biomass reservoir. The lysate is transferred to the purification system by the crew. The purification system uses a separate pump (1) and pinch-valve module (2) to load lysate into a crude lysate column (13). The column is washed with buffer (9), and recombinant protein is eluted into the product reservoir (11).

Design 3 uses an identical dewatering and lysis system as in design 2, but with a different protein purification system. Crew will transfer the lysate to the lysate chamber (14) of a continuous loop affinity membrane purification system (15). Rollers (17) move the affinity membrane through buffer chambers for protein binding (14), wash (9), and elution (10). A second wash chamber equilibrates the membrane for multiple cycles of protein binding. The crew transfers the eluted protein to a product reservoir (11).

### Comparative analysis of designs

Each design integrates different methods to complete the same bioprocessing steps (Fig. 5b). Multiple systems parameters including ESM, SCM, TRL, and degree of crew assistance were used to assess the three designs. SCM estimates complexity of life support systems by summing all the components and proposed interconnections of a specific design^16^. Larger SCM values are interpreted as more complex systems with potentially lower reliability. Figure 6 depicts the major components and interconnections used to calculate SCM values for each of the three designs. We also analyzed the degree of crew assistance because crew time was not incorporated into ESM calculations. Crew-assisted steps are highlighted in red or pink in Figures 5 and 6. TRL assesses technology maturity using a 9-point scale with TRL 9 being spaceflight-proven technologies^26^.

**Fig. 6.**
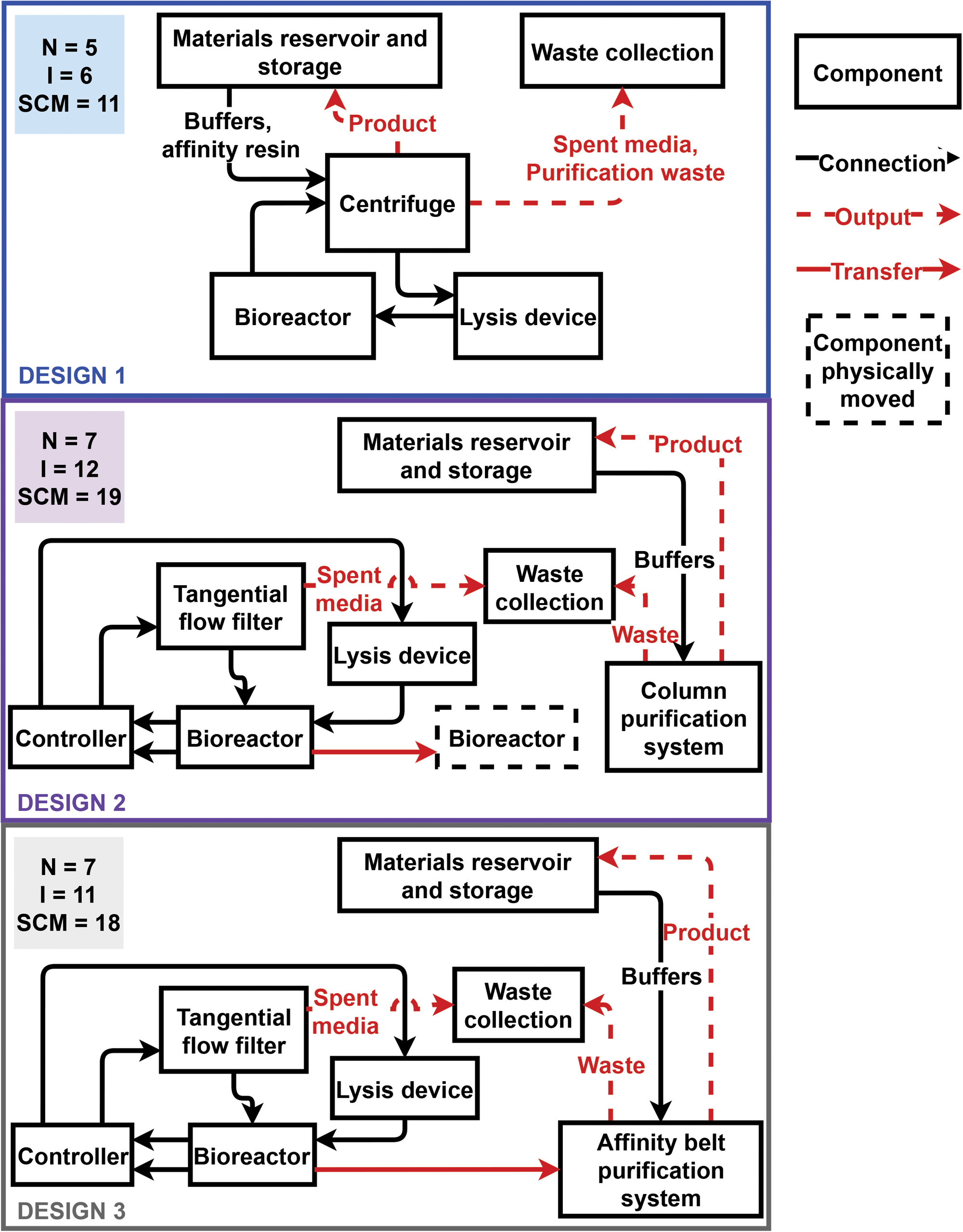
Top-level block diagram of the three bioprocessing designs used to calculate SCM. Major components (N) are in boxes. Individual interconnections (I) are diagrammed with arrows indicating the direction of material flow. Dashed arrows indicate outputs of the bioprocessing systems, and red arrows require crew assistance to complete the actions for the interconnection.

Figure 7 compares ESM, SCM, and degree of crew assistance for the integrated designs, while Table 4 reports TRL for the dewatering, cell lysis, and protein purification steps (Supplementary data 2). ESM was comparable for all three designs. Design 1 had the lowest SCM due to the multifunctional commercial countercurrent centrifuge that integrates dewatering, cell lysis, and protein purification. The ease of integration and automation in design 1 is also reflected in the low degree of crew assistance and higher TRL for dewatering and purification compared to the other designs. Although these metrics suggest design 1 will be more practical to implement, the differences in the metrics between the designs were not large enough to eliminate specific methods without experimental testing.

**Fig. 7.**
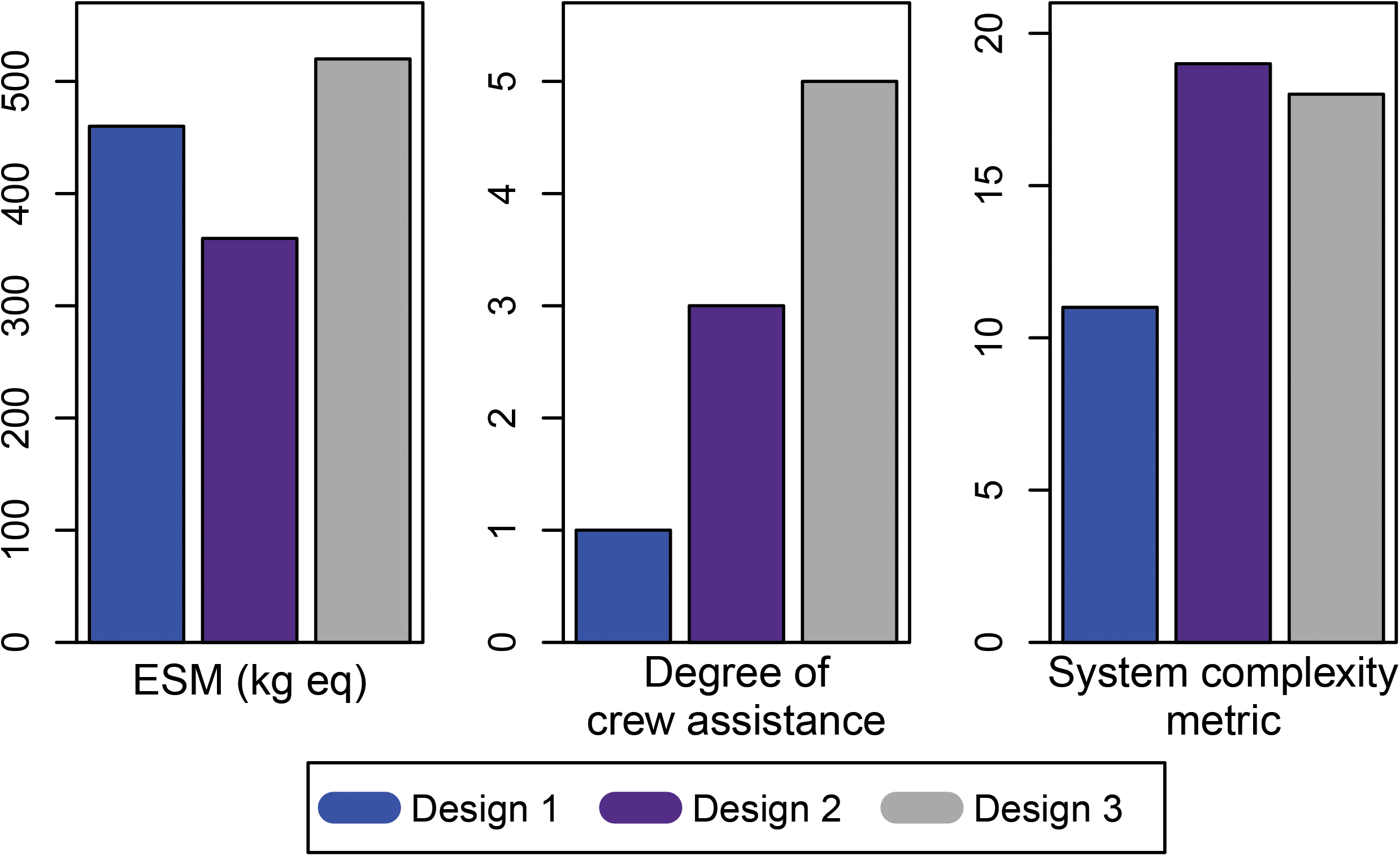
Design metrics for the three integrated bioprocessing designs. See Table 2 for total ESM values and Supplementary data 2 for detailed ESM calculations.

**Table 4.** TRL for each bioprocessing step of the integrated designs

| **Table 4. TRL for each bioprocessing step of the integrated designs** |  |  |  |
| --- | --- | --- | --- |
| **Step** | **Design 1** | **Design 2** | **Design 3** |
| Dewatering | 4 | 3 | 3 |
| Lysis | 9 | 9 | 9 |
| Protein purification | 3 | 3 | 2 |

## Discussion

Carbonic anhydrase improves the efficiency of liquid amine CO_2_ capture systems^5–7^, which is a candidate technology for deep space missions^1,2^. Application of these liquid amine systems requires a steady supply of carbonic anhydrase that may be met by long term storage, resupply or in situ bioproduction. Long term storage of active enzyme requires low temperature storage conditions, and frequent resupply is not possible for surface missions to Mars.

Bioprocessing system designs for space applications need to optimize mass, efficiency, power consumption, complexity, reliability, and ease of operation^37,38^. The initial space design process largely focuses on ESM to ascertain feasibility either in comparison to existing state of the art or other comparable technologies^12,14,25,39–44^. In this study, we employed ESM along with processing duration, SCM, TRL, and degree of crew assistance metrics to estimate reliability, potential for implementation, and ease of operation. Although carbonic anhydrase production was the use case scenario, the designs would be useful to purify any soluble recombinant protein with an affinity tag from a microbial chassis organism.

ESM comparisons were able to eliminate batch centrifugation and sonication as methods for integrated designs. However, the integrated bioprocessing designs have a narrow range of 357-522 kg eq ESM. Given these differences are not large enough for system selection, this metric cannot be used to prioritize designs for implementation. The ESM estimates are comparable to a proposed monoclonal antibody production system at 300-1,700 kg eq^45^ as well as a plant biomass system for food production at 300-1,300 kg eq per kg^10^. For theoretical designs such as the proposed biomanufacturing systems, a difference of less than 10-fold suggests similar infrastructure costs. On a mission-wide basis, the recombinant protein bioprocessing system may be reasonable for a Mars mission. The bioprocessing system adds only 2-3% to the physio-chemical-biological life support system proposed by ESA for a full Mars mission (∼18,000 kg eq)^46^.

Crew time and system complexity are important factors in space system design ^16,47^. Design 1 ranked best for number of crew assisted steps and SCM, which primarily reflects the level of integration for automated fluid handling in the commercial counterflow centrifuge. Additional engineering to integrate pumps and controllers would reduce both SCM and crew-assisted steps for designs 2 and 3. Design 1 also has higher TRL components indicating this design may be easier to implement for space, but a limitation of the study is a lack of empirical validation. All components were assumed to operate as intended and to be compatible within a design and many components have not been tested for the specific use case of recombinant protein purification. For example, the counterflow centrifuge in design 1 has not been used for bioprocessing of microbes. Empirical tests are needed to determine the relative efficiency of the selected technologies when integrated.

Although closed-loop systems and reusability are attractive long-term goals in space, they require increased crew time and system reliability^38,48,49^. On Earth, biomanufacturing often employs single-use technology for various reasons including improved safety as well as reduced contamination, footprint, and cost by eliminating cleaning and sterilization of components that contact biological material^50,51^. Single-use materials for bioprocessing is predicted to reduce crew time and to increase reliability. Consequently, the proposed bioprocessing designs use disposable components in the ESM calculations.

We also identified several factors that could influence a recombinant protein bioprocessing design that were difficult to quantify. For example, there are safety and shelf-life concerns for commercial affinity purification products. Affinity resins are stored in ethanol to prevent biological contamination. Ethanol interferes with the current Environmental Control and Life Support Systems on the ISS, creating a safety risk when large amounts of ethanol are in the cabin air or wastewater. The ethanol risk could be mitigated with an affinity membrane that is stored dry. Both commercial resins and affinity membranes have 1-2 year shelf lives, while a Mars mission is expected to require a 5-year shelf life for all systems^27^. Removing incompatible chemicals and extending the shelf life of the affinity matrix are both critical to advance space biomanufacturing of recombinant proteins.

It is possible that using different strategies to express recombinant proteins could reduce complexity and system mass. For instance, an affinity resin “teabag” can purify proteins secreted into the media without dewatering or lysis steps^52^. Although this strategy would simplify bioprocessing, it limits production to recombinant proteins that have activity after secretion. Moreover, E. coli secreted proteins are targeted to the periplasmic space and require an osmotic shock step to release protein into the media. Adding an osmotic shock still requires dewatering to concentrate biomass and the osmotic shock step would replace the cell lysis step.

This study illustrates multiple approaches to rank technologies for design of new integrated systems intended for space applications. The extreme environment of space, high cost of launching materials, and limited crew time drive systems to be automated and to minimize mass, power, and volume. Further development of space bioprocessing systems is expected to identify technologies that can be integrated and automated reliably, which could be translated to improved efficiency for industrial biomanufacturing on Earth.

## Methods

The operational scenario investigated was a six person crew on a 3-year Mars mission with 600 days of surface operations^31^. A total of 14.4 g of recombinant carbonic anhydrase was estimated to be required to maintain a liquid amine CO_2_ scrubber with active enzyme throughout surface operations. Affinity purification of the enzyme was assumed to be needed to reduce liquid amine side reactions with cellular debris. Sufficient carbonic anhydrase could be supplied by approximately 80 bioprocessing runs using 1 L cultures with a yield of 180 mg purified enzyme per run. Supplies and power for a total of 100 production runs were assumed to give sufficient redundancy for production runs that failed to meet expected yield.

A review of recombinant protein purification from unicellular microorganisms identified sub-process steps (Fig. 1). Essential steps for the operational scenario, such as lysis or purification, were retained. Biomass dewatering was investigated as an optional step. Non-essential steps, such as storage and drying, were eliminated to simplify potential workflows. A trade study of potential methods for each sub-process step was conducted, and ESM was used to evaluate individual technologies for in-space application. Technologies with comparable ESM were used to develop three designs, which were analyzed using four metrics described below.

### ESM metric

ESM was calculated using Equation 1 and the equivalency factors in Table 1 based on guidelines from Levri, et al. ^14^. Parameters for volume (V^ESM^), power (P^ESM^), thermal cooling (C^ESM^), cold storage (CS^ESM^), water (W^ESM^), and waste storage (WS^ESM^) used equivalency factors estimated for a nominal crewed surface mission to Mars^31^. CS^ESM^ assumed that the mass requiring cold storage had a density of 1000 kg/m^3^ and was stored in an freezer analogous to that currently used on the International Space Station (ISS)^31^. W^ESM^ and WS^ESM^ values were derived from a prior Mars mission scenario where a crew of six used 30 kg of water per crew member per day, and produced 1.5 kg solid waste per crew member per day^31,55^. W^ESM^ was based on a water processor similar to the ISS water treatment system and WS^ESM^ assumed that waste was stored within the habitat^55^. Crew time was excluded due to the uncertainties in estimating the practical level of automation that could be achieved after integrating technologies from different commercial manufacturers. Detailed assumptions and calculations of ESM are given in Supplementary data 1 and 2. ESM penalties from both 2015 and 2022 Baseline Values and Assumptions Documents were used for comparative calculations shown in Table 2^31,32^.

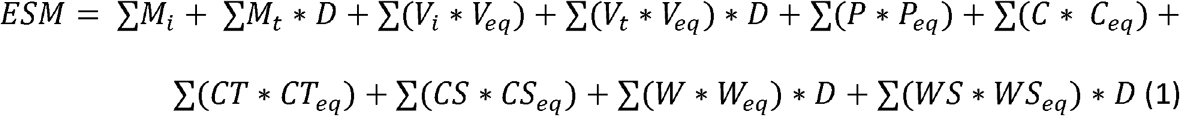

Where,

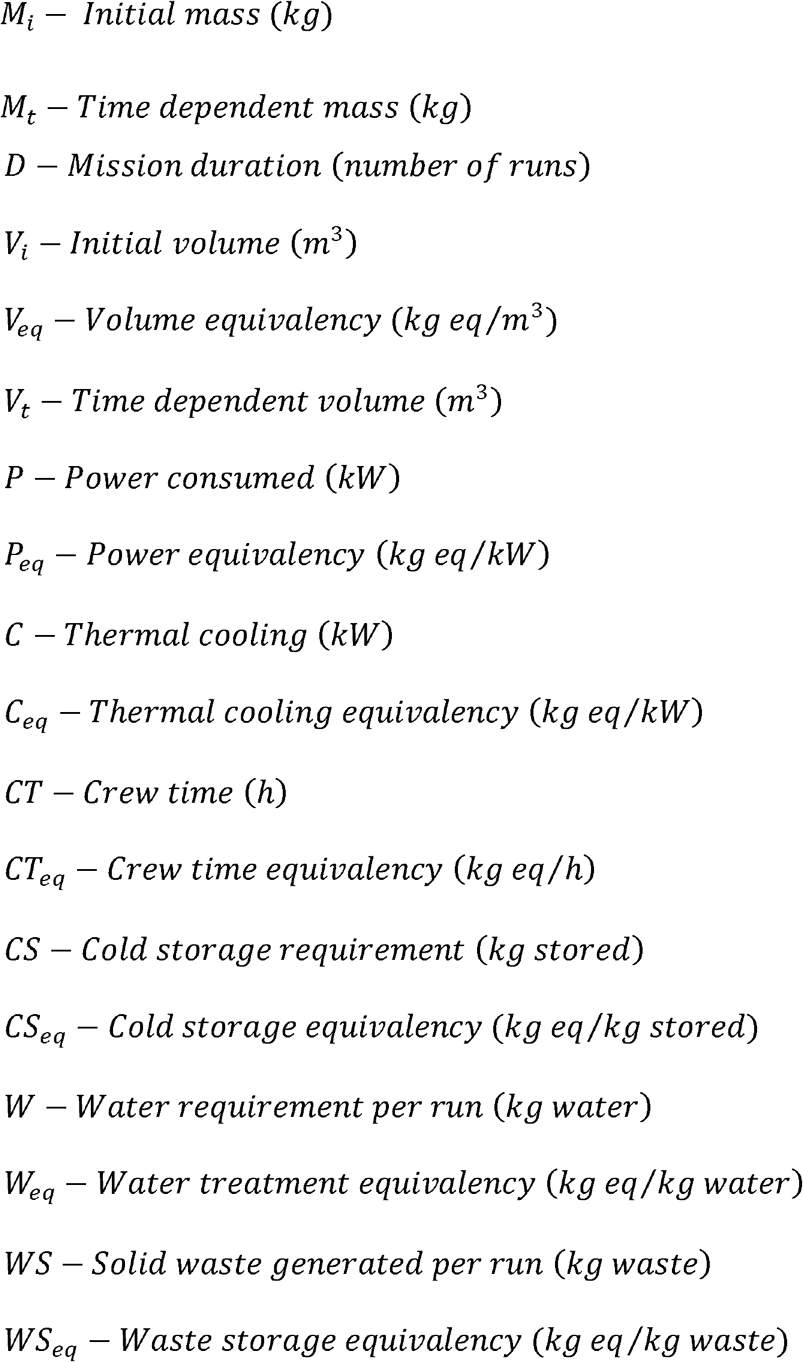

### SCM metric

Equation 2 gives the SCM calculation based on the number of components (N) and the number of one-way interactions (I) between components^16^.

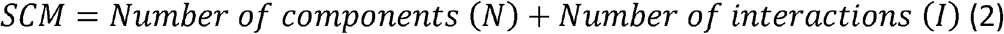

Components are integrated subsystems that perform one or more bioprocessing steps, such as equipment available from commercial vendors. Auxiliary parts such as valves, ordinary filters, and sensors were excluded with the rationale that integration of major components account for most of the system complexity, cost, and failure modes^16^. The number of one-way interactions (I) is derived from a top-level bioprocessing system block diagram based on physical connections between components (Fig. 6).

### TRL metric

A TRL was assigned for each of the dewatering, cell lysis, and purification options included in the three designs using existing NASA guidelines^26^.

### Degree of crew assistance metric

The degree of crew assistance was estimated based on whether the commercially available components were integrated and easily automatable. A crew support step was assumed if commercial technology was not readily available to automate a specific step in the proposed workflow of each design. The sum of crew assisted steps was the degree of crew assistance for the design.

### Dewatering technologies

Tangential flow filtration, batch centrifugation, and countercurrent centrifugation were three dewatering methods compared for this analysis. The dimensions and mass of the tangential flow filter were based on Xampler cartridge with 1 mm fiber diameter, 500 kDa pore size and 30 cm path length with a 3M housing (Cytiva, Marlborough, MA, USA). To control fluid flow through the tangential flow filter, an Ismatec Reglo ICC Digital Pump with 4-Channels and 8-Rollers (Cole-Palmer, Vernon Hills, IL, USA) was included in the ESM calculations. A modified version of the Drucker model 755VES swinging bucket centrifuge (Drucker Diagnostics, Port Matilda, PA, USA) has been approved for operation on the ISS, and the specifications of the commercially available centrifuge were used to calculate the ESM. Twenty disposable 50 mL tubes were assumed to harvest the 1 L culture volume during every run. The countercurrent centrifuge data was based on specifications for the CTS Rotea Counterflow Centrifugation System (Thermo Fisher Scientific, Waltham, MA, USA).

### Lysis technologies

The methods considered for lysis were bead beating, enzymatic lysis, a flow-through sonicator, and a large-volume probe sonicator. The bead beater analysis was based on the Claremont Biosolutions LLC (Upland, CA, USA) OmniLyse HL beadbeater flow-through lysis device. Although originally developed for small volumes, the OmniLyse HL unit was assumed to lyse large volumes with equal efficiency using extended processing times. The enzyme lysis protocol assumed that the biomass was incubated for 30 minutes at ambient temperature with 0.25 mg/mL lysozyme, 0.1 mL/mL Pierce universal nuclease (Thermo Fisher Scientific, Waltham, MA, USA), and 0.1% w/v Triton X at final concentration. Mass and volumes for ESM were calculated assuming the solid reagents had a density equal to NaCl (2.17 g/cm^3^), and liquid reagents had a density equal to water (1 g/cm^3^). The QSonica (Newton, CT, USA) ultrasound generator was modeled assuming a 1-inch replaceable tip for a large volume batch method and a Q500 FloCell unit for the flow through method.

### Protein purification technologies

Five His-tag affinity purification methods were compared using public domain product information available from commercial vendors. The Sigma Aldrich (St Luis, MO, USA) His-Select Ni affinity gel was used to represent affinity columns requiring clarified lysates. The Cytiva (Marlborough, MA, USA) HisTrap FF was an exemplar of a crude lysate column that does not require clarification before sample loading. Batch resin purification specifications were estimated for the resin “teabag” method from Castaldo, et. al.^52^. Millipore Sigma (Burlington, MA) HIS-Select® Nickel Magnetic Agarose Beads specifications were used to estimate the amount of resin required for batch purification with magnetic beads, while mass and volume of the beads were assumed to be equivalent to affinity resin. For affinity membrane purification, Capturem large volume filters (Takara Bio USA Inc., San Jose, CA, USA) were used to estimate the quantity of membrane required, while mass and volume of the membrane was assumed to be equivalent to Whatman filter paper (200 g/cm^2^).

The buffer composition was 20 mM HEPES, 500 mM NaCl and 20 mM imidazole for the binding buffer, and 20 mM HEPES, 500 mM NaCl and 200 mM imidazole for the elution buffer for all the methods. Buffer volumes were modeled using manufacturer protocols or the resin “teabag” method^52^.

### Bioprocessing system designs

All three post-growth bioprocessing system designs assumed a Claremont Biosolutions LLC (Upland, CA, USA) OmniLyse® HL disposable bead beater for in-line cell lysis from the biomass reservoir. The first design included a Thermo Fisher Scientific (Waltham, MA, USA) CTS Rotea Counterflow Centrifugation System with disposable, single use kits for processing. This counterflow centrifuge includes integrated pinch-valves, a peristaltic pump, and a controller for automation. QIAexpressionist Ni-NTA resin (Qiagen Inc., Valencia, CA, USA) was included for affinity purification.

The second design uses a MidGee ultrafiltration cartridge UFP-5-C-MM01A (Cytiva, Marlborough, MA, USA) and an Applikon Biotechnology (JG Delft, Netherlands) “my-control” with built in peristaltic pumps for biomass dewatering using tangential flow filtration and subsequent cell lysis. For protein purification, an Automate Scientific (Berkeley, CA, USA) Perfusion System and ValveLink8.2 Perfusion Controller were assumed to integrate with an Masterflex (Gelsenkirchen, Germany) L/S® Digital Drive peristaltic pump with an Easy-Load® 3 Pump Head for Precision Tubing. This fluid control system was assumed to automate affinity purification in a Cytiva 5 mL HisTrap FF crude lysate column (Marlborough, MA, USA).

Dewatering and lysis components in design 3 were identical to design 2. Protein purification was based on a non-commercial affinity belt system^57–59^. Rollers were proposed to move an affinity membrane belt continuously through chambers containing the lysate, wash buffers, and elution buffers. The total volume of the system as assumed to be 0.005 m^3^ including the chambers, walls, and rollers. The mass was assumed to be 500 g. One Transmotec Inc. (Burlington, MA, USA) 12 V, 2A DC motor was included to operate the rollers.

Disposable materials such as bags, and sterile filters for all three designs were estimated for the system based on commercial options and material properties, while the additional mass of tubing and luers were assumed to be 0.1 kg for all designs.

## Data Availability

All source data and ESM calculations needed to replicate the study are provided in the manuscript and Supplementary data 1 and 2.

## Acknowledgements

We thank Aditya Hindupur for discussions in the initial phase of the study. This work was funded by the NASA Space Technology Mission Directorate Game Changing Development Program as part of the Space Synthetic Biology Project.

## Author Contributions

M.S. and M.B.P. completed the trade study investigation, formal analysis, and wrote the manuscript. M.D. completed data collection, formal analysis, and edited the manuscript. H.W.J. provided systems engineering expertise for formal analysis and edited the manuscript. J.A.H. conceived and supervised the project and edited the manuscript. F.M.D., J.M.G., and A.M.S. contributed to methods development, validation, and visualization of the results, and edited the manuscript.

